# A novel imaging biomarker for survival prediction in EGFR-mutated NSCLC patients treated with TKI

**DOI:** 10.1101/681577

**Authors:** Annabelle Collin, Vladimir Groza, Louise Missenard, François Chomy, Thierry Colin, Jean Palussière, Olivier Saut

## Abstract

EGFR-mutated non-small cells lung carcinoma are treated with Tyrosine Kinase Inhibitors (TKI). Very often, the disease is only responding for a while before relapsing. TKI efficacy in the long run is therefore challenging to evaluate. Our objective is to derive a new imaging biomarker that could offer better insights on the disease response to treatment. This study includes 17 patients diagnosed as EGFR-mutated non-small cell lung cancer and exposed to an EGFR-targeting TKI. The early response to treatment is evaluated with 3 computed tomography (CT) scans of the primitive tumor (one before the TKI introduction and two after). Using our knowledge of the disease, an imaging biomarker based on the tumor heterogeneity evolution between the first and the third exams is defined and computed using a novel mathematical model calibrated on patient data. Defining the overall survival as the time between the introduction of the TKI treatment and the patient death, we obtain a statistically significant correlation between the overall survival and our imaging marker (p = 0.009). Using the ROC curve, the patients are separated into two populations hence the comparison of the survival curves is statistically significant (p = 0.025). Initial state of the tumor seems to have a role for the prognosis of the response to TKI treatment. More precisely, the imaging marker - defined using only the CT scan before the TKI introduction - allows us to determine a first classification of the population which is refined over time using the imaging marker as more CT scans become available. This exploratory study leads us to think that it is possible to obtain a survival assessment using only few CT scans of the primary tumor.

## Introduction

Tyrosine Kinase Inhibitors (TKI) were shown to be effective in the treatment of EGFR-mutated non-small cell lung carcinoma (NSCLC) [1]. They are currently used as first-line treatment for patients of stage IV. The EGFR gene corresponds to the Epidermal Growth Factor receptor, which belongs to the family of receptors with tyrosine kinase activity. The alteration of the EGFR gene in lung cancer occurs in 5 to 30% of cases, depending on the patient origin (10% of Caucasian patients, 40% of non-smoking Caucasian patients and up to 60% of non-smoking Asian patients), see [2]. TKI treatment may be successful for a time but their efficacy in the long run is challenging to evaluate. For example, in [3], the authors estimate the relapse median time at 10 months.

Latest advances in oncology and the discovery of many different sub-types of cancer, partly because of genomic alterations, open the way to a personalized medicine [4]. There is a need of new tools combining different types of available data to help to choose the best treatment for each patient. Medical imaging has an important role to play in this context as these patients are routinely monitored using CT scans. The most current used CT scan evaluation - in particular for lung cancers - is the RECIST (Response Evaluation Criteria In Solid Tumors) which consists in measuring the largest diameters of target lesions [5, 6]. Most recent studies have shown the interest of the tumor volume evaluation which is more precise and has a better reproducibility in particular concerning the evaluation of non-small cell lung cancers [7–9]. Concerning EGFR-mutated non-small cells lung carcinoma, previous works have studied the correlations between the initial reduction of the primary tumor and the overall survival. One can find contradictory results in the literature: in [10, 11], a significant correlation has been established but more recently in [12] this correlation has not be validated using another database. Many recent studies propose to use radiomic approaches which consist in extracting a large number of quantitative features from medical images using data-characterization algorithms. In non-small cell lung cancers, various tumor heterogeneity markers may be computed, see [13] for a proposal for harmonization of methodology. Then, they can be related for example to the distant metastasis probability [14]; to predict pathological response after neoadjuvant chemoradiation [15]; to indicate tumor response to radiation therapy [16]; to advance clinical decision-making by analyzing standard-of-care medical images [17] and to establish independent marker of survival time [18]. In [19], the authors even show that radiomics may help identify a general prognostic phenotype existing in both lung and head-and-neck cancer. These approaches have also been used for EGFR-mutated non-small cell lung cancer. For example, radiomic approaches may predict EGFR mutation status without requiring repeated biopsy acquisitions [20, 21] but also identify tumor heterogeneity markers which can be related to early EGFR TKI failure [22].

In this study, instead of testing classical heterogeneity markers, we introduce a novel imaging biomarker that quantifies the evolution of the heterogeneity of the primitive tumor of patients with EGFR-mutated non-small cells lung cancers over time. This criterion is defined using our knowledge of the disease and can be biologically interpreted. The objective of this work is to study the value of this novel marker to predict overall survival in order to help clinicians to detect EGFR TKI treatment failure earlier.

## Materials and methods

### Patients

A monocentric retrospective cohort study has been conducted on patients with a biopsy-proven non-small-cell lung carcinoma - presenting an identified or suspected EGFR (epidermal growth factor receptor) mutation, (established by a TKI clinical benefit of more than 6 months) - which are non-accessible to local treatment (stage IIIB or IV). Patients were included in the study if 3 CT-scans were available: one before the first introduction of TKI treatment and two after. The study was approved by Institut Bergonié and IRB approval was obtained for use of the CT images. Informed consents of data collection were waived for research from each patient, in accordance with the related policy of Institut Bergonié.

### Treatment

All patients were exposed to an EGFR-targeting TKI. Two molecules were used: gefitinib (IRESSA®, Astra-Zeneca) and erlotinib (TARCEVA®, Roche). These two therapies were given until progression, unacceptable toxicity, patient refusal to continue treatment or death.

### Imaging and biomarkers

Evaluation scans were done every 2 to 6 months. The acquisition was performed after an injection of iodized contrast agent at portal phase on the thorax, the abdomen and the pelvis and then at late-arterial phase on the encephalon. We consider 3 CT-scans (given at times *t*_0_, *t*_1_ and *t*_2_). The first one is acquired immediately before TKI treatment and the second and the third ones are the 2 first ones after the first introduction of TKI treatment. Time *t*_0_ is the baseline. Tumors were delineated using a semi-automatic segmentation library that relies on a deformable model [23]. All the delineations were validated by a junior and a senior radiologists.

Using these images, we derive various biomarkers. Defining by *V* (*t*) the volume of the tumor at time *t*, we define the following set of biomarkers (computed from the volume):

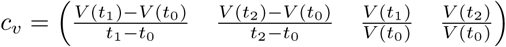

corresponding respectively to the initial slope of volume decreasing (between *t*_0_ and *t*_1_), to the slope of volume decreasing (between *t*_0_ and *t*_2_), to the initial percentage of volume decreasing (between *t*_0_ and *t*_1_) and to the percentage of volume decreasing (between *t*_0_ and *t*_2_).

On CT, high intensities (lighter colors on the image) correspond to high tissue densities and therefore to high cellularity and proliferation while intensities around 0 correspond to water and necrosis. We therefore split the set of voxels of the images into two classes. The first class contains the voxels whose values are non-negative while the second class is formed by non-positive intensities voxels. We will refer to the first class (with positive intensities) as being the proliferative one while the second one will be referred as the necrotic one. We denote by *P* (*t*) the volume of the set of voxels whose intensities are positive within the tumor on the exam at time *t*. We then compute the ratio of this proliferative-like compartment with respect to the total volume 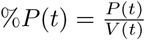. We define the following set of biomarkers (based on the heterogeneity):

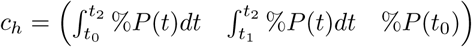

corresponding respectively to the area under the curve (AUC) of the quantity %*P* (*t*) between *t*_0_ and *t*_2_, to the AUC of the quantity %*P* (*t*) between *t*_1_ and *t*_2_, and to the initial value of the quantity %*P* (*t*).

Using the first 3 CT-scans, we extract *V* (*t*_0_), *V* (*t*_1_), *V* (*t*_2_), *P* (*t*_0_), *P* (*t*_1_) and *P* (*t*_2_) and approximate the imaging marker 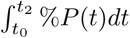 with the trapezoidal rule. However, this strategy has limitations. Indeed, the acquisition times *t*_0_, *t*_1_ and *t*_2_ and the CT image noise may introduce some instability in the computation, for example if *t*_1_ is too close to *t*_0_ or *t*_2_.

We therefore choose another strategy that consists in using a mechanistic model in order to fit the values *P* (*t*_0_), *P* (*t*_1_) and *P* (*t*_2_) continuously on the time interval [*t*_0_, *t*_2_]. This model relies on the evolutions of two populations of cells (proliferative cells and quiescent or necrotic compartment). This model is parametrized using the values *V* (*t*_0_), *V* (*t*_1_), *V* (*t*_2_), *P* (*t*_0_), *P* (*t*_1_) and *P* (*t*_2_) defined above. It then provides an evaluation of *P* (*t*) and *V* (*t*) at any time between *t*_0_ and *t*_2_, see Fig. 1. The model and its personalization are presented in the Supplementary Materials. On Fig. 1, the patient had a CT-scan at 109 days and at 206 days. The x-axis represents the time in days, while the volume is reported on the y-axis. The blue points denote the measured volumes on exams while the red and green points show respectively the volume of proliferative and necrotic compartments. The blue (resp. red, green) curve describes the evolution of the volume (resp. density of proliferative cells, density of necrotic cells) given by the mechanistic model that fits with the data. The purple curve gives the evolution of the ratio of the proliferative compartment with respect to the volume. Figure 8 (see Supplementary materials) gives the curves for all the patients.

**Fig 1.**
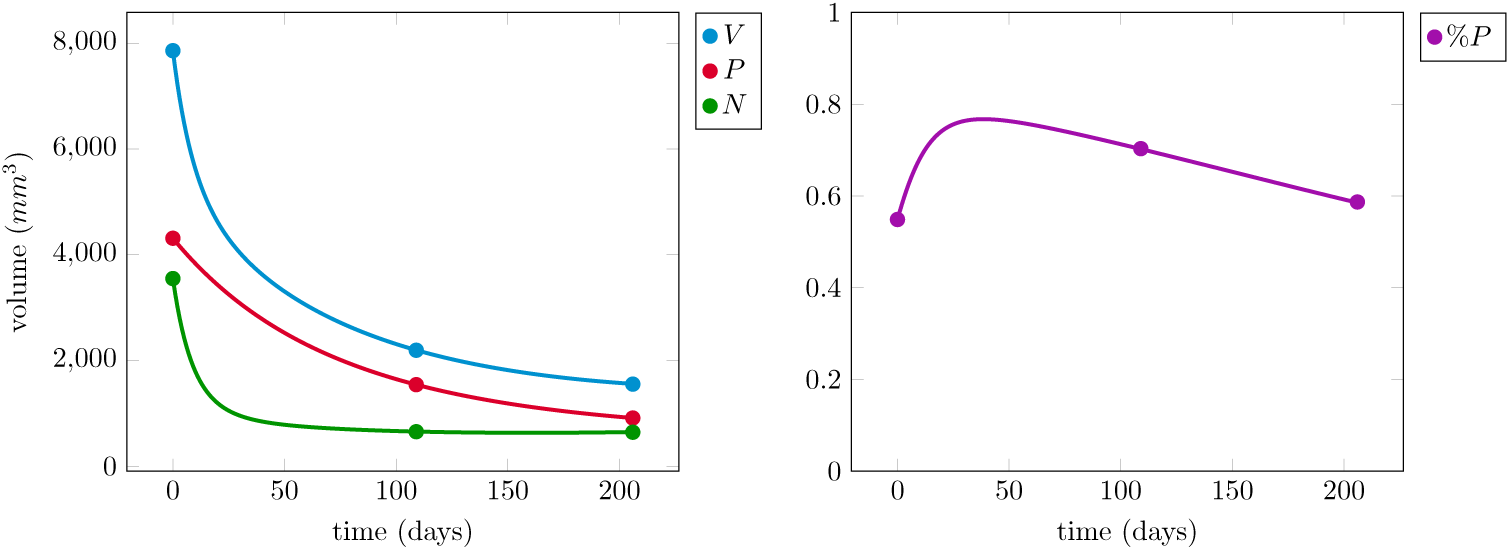
Left - Evolution of the densities of proliferative (red) and necrotic (green) cells for 1 patient with the evolution of the volume (in blue). Right – Evolution of the ratio of the proliferative compartment.

### Statistics

Whenever appropriate, standard statistics are presented as mean±standard-deviation and number (percentage). We define the overall survival (OS) as the time between the introduction of the TKI treatment and the patient death. Survival curves were computed using the Kaplan-Meier estimator and compared using Log-Rank tests. The association of survival failure with each investigated biomarker was tested using Cox regression. Prediction performances of the biomarkers were assessed using ROC curves. The appropriate statistical tests were performed when required with a significance threshold set to p = 0.05. The mechanistic model was fitted using the Monte Carlo method. All computations were performed using `Matlab-R2015a`.

## Results

A population of 25 patients has been collected at Institut Bergonié (Bordeaux, France) between 2006 and 2013. We have kept 17 patients among these 25 patients. We have excluded 2 cases for which the CT-scan before the TKI introduction was not available and 6 cases for which it was not possible to delineate the tumor (miliary disease, patients without any discernable lesion e.g. with pleural effusion or atelectasis). Table 1 presents the patient cohort: age, sex, smoking information, mutation, stage and if the patient had a treatment before the TKI introduction.

**Table 1.** Presentation of the patient cohort: sex, age, smoking, mutation, stage, previous treatment before the TKI introduction.

| Characteristics |  |  |
| --- | --- | --- |
| Sex | Women | 14 (82 %) |
|  | Men | 3 (18 %) |
| Age |  | 65 ± 11 |
| Smoking | Yes | 1 (6 %) |
|  | No | 10 (59 %) |
|  | Unknown | 6 (35 %) |
| Mutation | Exon 19 | 17 (41 %) |
|  | Exon 21 | 7 (41 %) |
|  | Exon 18 | 1 (6 %) |
|  | Unknown | 2 (12 %) |
| Stage | IV | 16 (94 %) |
|  | IIIB | 1 (6 %) |
| Previous treat. | No | 11 (65 %) |
|  | Yes | 6 (35 %) |

First of all, there is no significant correlation between each volume-based biomarker gathered in the vector

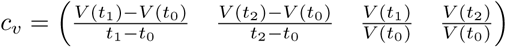

and the overall survival (p = 0.48, p = 0.36, p = 0.23 and p = 0.17).

We will now focus on the heterogeneity-based biomarkers gathered in the vector

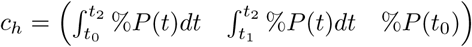

The values of the first heterogeneity-based biomarkers

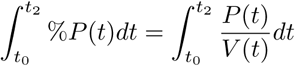

for all the patients are plotted in Fig. 2.

**Fig 2.**
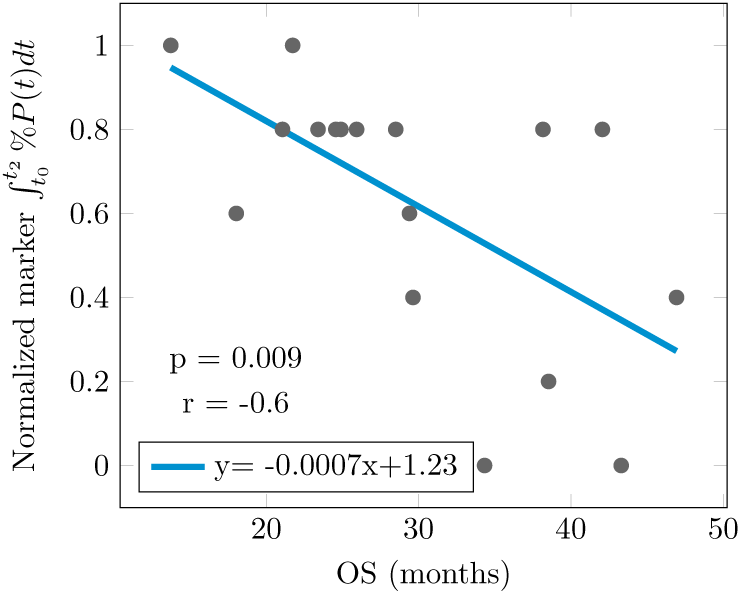
Imaging marker 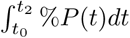 after normalization (y-axis) versus overall survival in months (x-axis) for each patient. Linear regression in blue.

The values of the criteria based on the evolution of the tumor heterogeneity have been normalized using its maximum value and is reported on the y-axis. The x-axis corresponds to the value of the overall survival. The blue curve corresponds to the linear regression. The correlation between the overall survival and the imaging marker is statistically significant (p = 0.009, r =-0.6). The population may clearly be divided into two populations. In particular, patients with a short survival time have a large value of the biomarker. In order to find the best threshold to classify the patients, we need to set a survival threshold. Based on the survival histogram of the 17 patients given in Fig. 3 and for clinical relevance, we set the survival threshold at 30 months.

**Fig 3.**
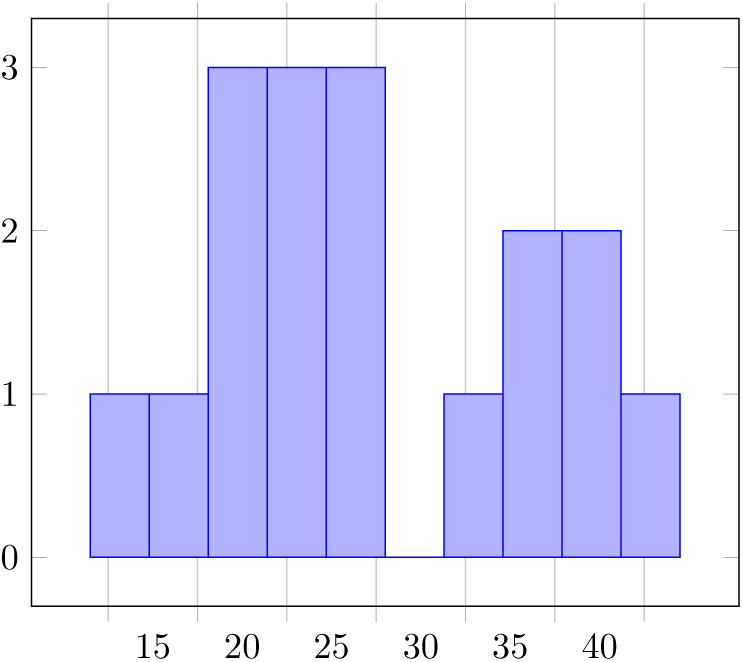
Survival histogram (17 patients).

The ROC curve (AUC = 0.81) is given in Fig. 4 (see blue curve). Using Fig. 4, we see that a good compromise consists in taking a normalized threshold for the biomarker of 0.4 that is optimal with a sensibility of 0.9 and a specificity of 0.7..

**Fig 4.**
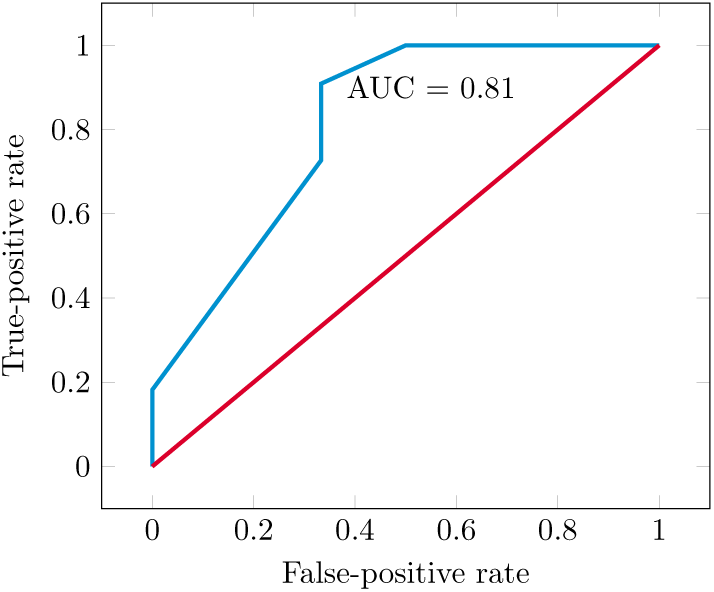
ROC curve of the imaging marker 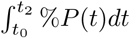 (after normalization).

Finally, survival curves are given in Fig. 5: in red the survival curve of the full population and in blue (resp. in green) the survival curve of the population with a short (resp. with a large) imaging marker. The normalized threshold value 0.4 obtained with the ROC curve is used to discriminate the patients. The comparison of the survival curves of these two populations is statistically significant (p = 0.025, hazard ratio = 0.25 with a 95% confidence interval equals to 0.09 - 0.7).

**Fig 5.**
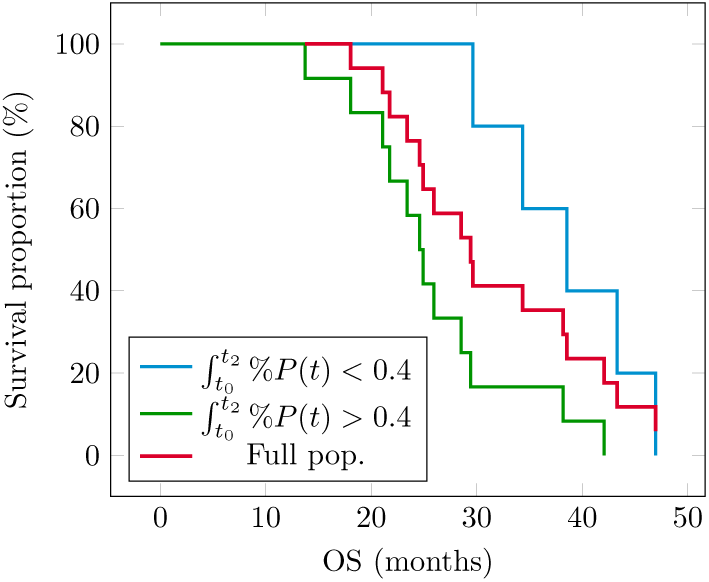
Survival curves (full population in red; patients with a small (resp. large) value of the imaging marker 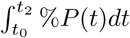 in blue (resp. in green).

However, the strategy is not efficient if we only use the time average between *t*_1_ and *t*_2_ to define the imaging biomarker. Indeed there is no correlation between the second heterogeneity-based biomarker 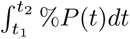 and the overall survival (p = 0.34). This means that the initial status of the tumor plays a major role in the evaluation of response to the TKI treatment. More precisely, there is a significant correlation between the third heterogeneity-based biomarker %*P* (*t*_0_) and the overall survival (p= 0.034) even if the biomarker based on the 3 CT-exams is six times more significant (p = 0.009), see Fig. 6. Concerning the survival curves, we obtain equivalent results (p = 0.036 instead of p = 0.025).

**Fig 6.**
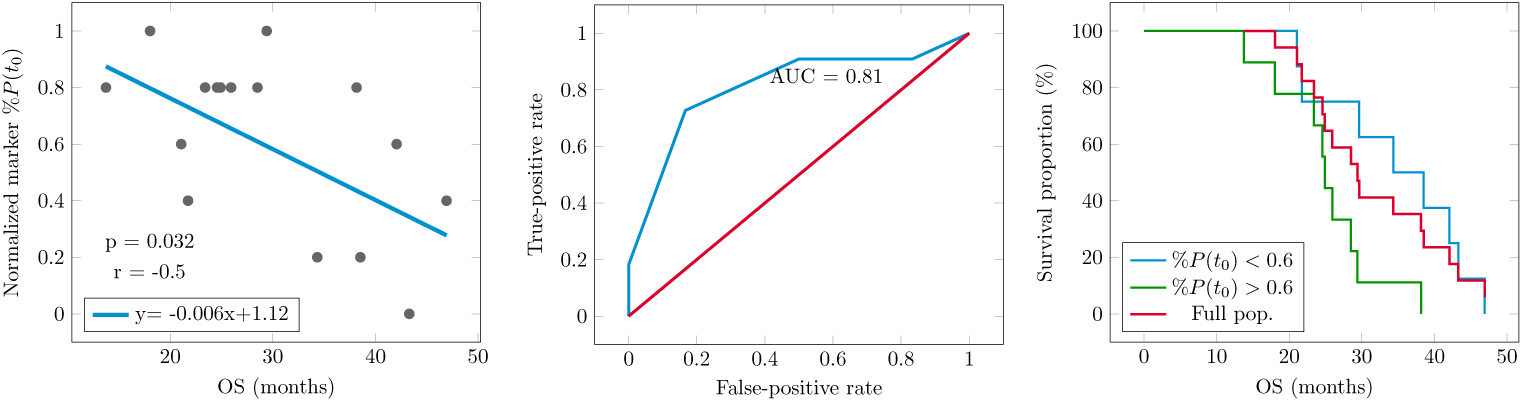
Left-imaging biomarker %*P* (*t*_0_) versus overall survival. Middle - ROC curve. Right – Survival curves (the threshold value has been fixed using the ROC curve).

## Discussion

In this work we have studied the ability of volume-based and heterogeneity-based imaging biomarkers to predict the survival in EGFR-mutated NSCLC patients treated with TKI.

Our first result is that there is no correlation between volume-based imaging biomarkers and survival. This finding is consistent with the work [12] who noticed a lack of association between tumor shrinkage and long-term survival. This illustrates why the response to TKI treatment is difficult to estimate using only the evolution of tumor volume (or RECIST).

Our second result is that heterogeneity-based imaging biomarkers may help predict short-term survival. More precisely, we propose an imaging biomarker 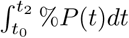 that is able to discriminate patients with short-time survival using only 3 successive CT-scans (an imaging monitoring performed as standard of care for such patients). To the best of our knowledge, this is the first study in which a mechanistic model based on disease knowledge has been used to predict the outcome in EGFR-mutated NSCLC patients treated with TKI. This shows that characterizing the content of the tumor and its dynamics using mathematical models might provide valuable information to guide clinical decisions.

The main strength of this work lies in the fact that the imaging marker is based on a mathematical description of the alleged underlying pathophysiological processes rather than purely empirical observations. As a consequence, the value of the biomarker may be given a phenomenological meaning, an interpretation that would be lacking otherwise. A large value of the biomarker means that the proportion of proliferative cells does not decrease over time (even if the volume of the lesion is decreasing). In the opposite case, a small value of the marker implies a decrease of the proportion of the proliferative compartment, even if the response is modest in terms of whole tumor size.

We show that the use of the first image (CT-scan acquired before TKI introduction) is of paramount importance for biomarker to predict the survival. More precisely, the imaging biomarker computed using only this first image provides a first classification of the patient that can be incrementally improved using the imaging marker as more follow-up CT-scans become available.

In addition to biomarkers, the mathematical model (see Supplementary Materials) offers more information on the mechanism of the response to treatment. The evolutions of the proliferative and necrotic compartments given for each patient might also be useful to personalize therapy, see Figure 8. In particular, it would be interesting to study the ability of this mathematical model to detect acquired resistances to TKI earlier, especially T790M mutation, a feature associated with bad prognostic [24].

An important limitation of this work is the small sample size and its retrospective nature. A second limitation concerns the assumptions of the mechanistic model: tumor heterogeneity as measured using CT-scan is assumed to be related to the proportion of proliferative and necrotic cells. Confirming this hypothesis would require a histological assessment of the whole tumor, which is possible only in patients who undergo surgery.

## Conclusion

We have shown that the initial volume (or RECIST) evolution under TKI is not sufficient to predict the survival while the tumor heterogeneity before the TKI introduction is a major prognosis factor and provides a first classification of patients. Furthermore, this first classification can be incrementally improved using the imaging marker that summarizes the early evolution of the tumor heterogeneity as soon as more CT scans are available. Short term perspectives of this work are about increasing the size of the cohort and improving the segmentation process in order to be able to include patients with non-delineated tumors.

## Acknowledgments

This study was supported by the French Laboratory of Excellence TRAIL ANR-10-LABX-57.

## Supplementary materials

### Presentation of the mathematical model

We denote by *P* (resp. by *N*) the volume of proliferative (resp. quiescent or necrotic) cells. We have *P* + *N* = *V* where *V* corresponds to the volume of the lesion. The evolution of the volume of proliferative cells is supposed to satisfy the following equation:

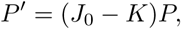

where the quantity *J*_0_ is the growth rate and we assume it constant in order to keep a model with identifiable parameters. The quantity *K* corresponds to the decreasing rate due to the TKI treatment. We assume that it follows a Gompertz-like law:

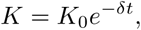

where *δ* is an unknown parameter and *K*_0_ the initial decreasing rate of *P*. We assume that when exposed to the treatment, the proliferative cells die and form the necrotic compartment. The evolution of the density *N* of this necrotic compartment is supposed to satisfy the following equation:

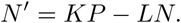

The quantity *L* is the evacuation rate of the necrotic compartment. We assume that it follows a Gompertz-like law:

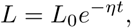

where *η* is an unknown parameter and *L*_0_ the initial evacuation rate of *N*.

This leads to the following ordinary differential system

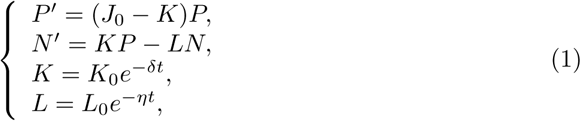

in which the quantity *P* can be explicitly determined:

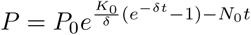

and the quantity *N* can be numerically approximated. Figure 7 illustrates this model in the formalism of compartment models.

**Fig 7.**
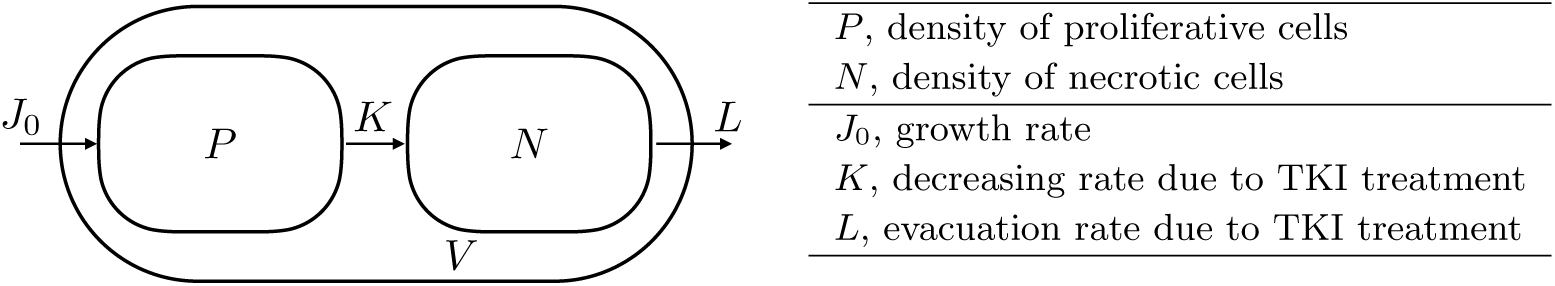
Modeling of temporal evolution of proliferative and necrotic cells.

**Remark 1** *Please note that this model may be derived from a spatial PDE model as follows. Let 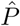 (resp. 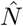) be the spatial density of proliferative (resp. necrotic or quiescent) cells and 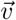, the velocity field that describes the evolution of the tumor over time. Following [25–27], the tumor can be described by the evolution in space and time of population of 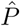 and 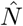*,

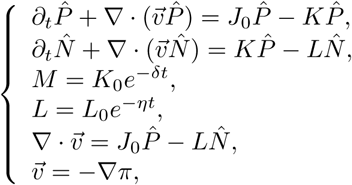

*where the last equation closes the system using a Darcy law with π the pressure. Using Reynolds theorem, this system of partial differential equation is related to System* (1) *by*

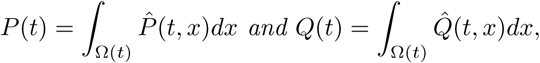

*where* Ω *is the tumor domain.*

### Model parametrization

The model has 5 parameters: *K*_0_, *δ, J*_0_, *L*_0_ and *η* that we want to estimate using (the subscript *d* is used to design the data):

- the volumes *V*_*d*_(*t*_1_) and *V*_*d*_(*t*_2_) (*V*_*d*_(*t*_0_) is used for the initial condition),
- the proliferative parts *P*_*d*_(*t*_1_) and *P*_*d*_(*t*_2_) (*P*_*d*_(*t*_0_) is used for the initial condition).

The parametrization is done into two steps. We start by estimating *K*_0_, *J*_0_ and *δ* by minimizing

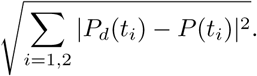

Two cases are possible:

- Case *P*_*d*_(*t*_0_) > *P*_*d*_(*t*_1_) > *P*_*d*_(*t*_2_): we assume that the density of proliferative cells is decreasing and we search the parameters as

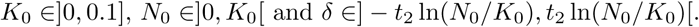
- Case *P*_*d*_(*t*_0_) > *P*_*d*_(*t*_1_) and *P*_*d*_(*t*_1_) < *P*_*d*_(*t*_2_): we search the parameters as:

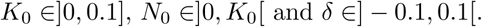

The set of parameters is not unique. However, if we consider two sets of parameters which give small errors (∼ 10^−3^), the variations of *P* are very close. The second step consists in estimating *L*_0_ and *η* by minimizing

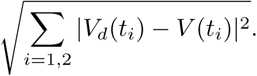

As we consider that the TKI treatment is still acting (for all the patients, we have *V*_*d*_(*t*_0_) > *V*_*d*_(*t*_1_) > *V*_*d*_(*t*_2_)), the sets of parameters for which *V* is not strictly decreasing are rejected.

## Results

The evolutions of the densities of proliferative (red) and necrotic (green) cells for the 17 patients with the evolution of the volume (in blue) are presented in Fig. 8.

**Fig 8.**
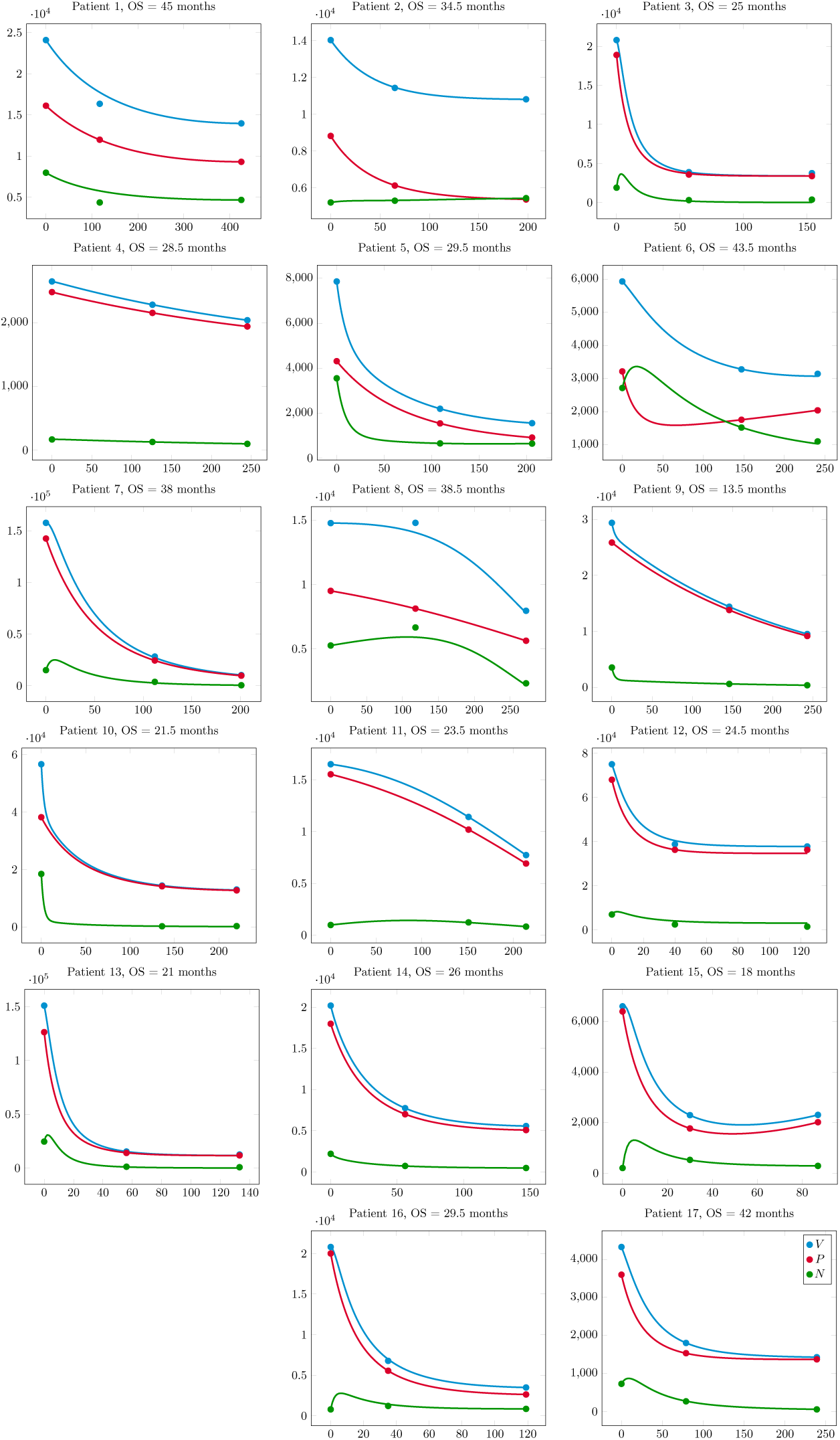
Evolutions of the densities of proliferative (red) and necrotic (green) cells for 17 patients with the evolution of the volume (in blue).

